# Striatal calcium transients detected by fiber photometry propagate to axons

**DOI:** 10.1101/2023.10.09.560813

**Authors:** David Matthew Lipton, Mohammad Tamimi, Itay Shalom, Tomer Sheinfeld, Ben Jerry Gonzales, Maya Groysman, Ami Citri

## Abstract

ARISING FROM: A. Legaria et al., Nature Neuroscience https://doi.org/10.1038/s41593-022-01152-z (2022). Calcium fiber photometry is a popular technique for recording the activity of neuronal populations defined by their gene expression or connectivity. In a recent study, Legaria et al., present evidence that the calcium signal recorded with fiber photometry primarily reports local fluctuations in neuropil Ca^2+^, rather than somatic Ca^2+^ influx corresponding to neural firing, as has been assumed by the field. This raises the question of whether fiber photometry transients are a valid measure of the propagation of information from neural soma to their axons. We addressed this question directly, recording coincident activity from both the somato-dendritic region and downstream axons of striatal neural populations. Our findings demonstrate that calcium events are reliably propagated to axons, supporting the interpretation that these events reflect neuronal firing.

---

A central goal of neuroscience is to decipher the neuronal activity patterns driving behavior. Different techniques have been developed to enable the recording of neural activity at varying levels of spatial and temporal resolution^1^. Within this suite of technologies, fiber photometry, when used with a Genetically Encoded Calcium Indicator (GECI) such as GCaMP^2^, is a useful technique for identifying correlations between Ca^2+^ flux in populations of neurons defined by their genetics or connectivity and the behavior of a recorded subject^1,3–6^. Calcium activity is thought to be a proxy of neuronal firing in the vast majority of neurons^7^: When a neuron fires action potentials, the voltage changes in the soma cause voltage-dependent Ca^2+^ channels to open, and the somatic Ca^2+^ concentration to increase, generating an increased GCaMP signal. Yet, other biological events can cause Ca^2+^ flux and subsequent GCaMP fluorescence changes as well^7^. For instance, dendritic potentials are also characterized by increased dendritic Ca^2+^ and subsequent GCaMP activity^7,8^. When recording Ca^2+^ flux with fiber photometry, due to the fact that all of the recorded fluorescence is combined together into a single measurement it is not possible to differentiate these different sources of Ca^2+^. Spatially, both somatic Ca^2+^ and neuropil Ca^2+^ (mostly dendritic by volume) are combined together to generate a composite GCaMP fluorescence signal recorded at the fiber tip^9^, raising the question of the relative contributions of each of these two sources of fiber photometry signal. This issue was recently brought to the attention of the field by Legaria et al., Nature Neuroscience 2022^8^. In this report, it is emphasized that this issue is particularly relevant for neurons that have widely arborizing dendrites, such as striatal Spiny Projection Neurons (SPNs).

We have been studying the activity of neurons in the ventrolateral aspect of the dorsal striatum (VLS)^10,11^, a striatal region involved in orofacial behavior^12,13^, and have recently utilized fiber photometry to address the relationship between neural activity in populations of VLS neurons and orofacial behavior. The concern raised by Legaria et al., made us re-evaluate this approach: If the photometry signal does not reliably reflect neuronal firing, but rather dendritic integration that is uncoupled to action potentials, then it dramatically changes the interpretation of our recorded photometry signal. We therefore sought to develop a direct strategy to judge the extent to which the signals we observe with fiber photometry from the striatal SPNs report the firing of action-potentials, which subsequently propagate to axons.

To this end, we injected the VLS of A2a-Cre mice with AAV9-diO-GCaMP6s and implanted fibers bilaterally in both the VLS and in the GPe, where the axons of VLS neurons project. The VLS-implanted fiber can collect GCaMP signals from both the soma region and neuropil, while the GPe fiber records Ca2+ flux exclusively from the downstream axons (Fig. 1A). We could thus simultaneously record the fiber photometry signals from VLS indirect-pathway SPN neurons (iSPNs) in the VLS and their axons in the GPe and then analyze the degree to which these two signals align. Post-experiment histology showed GCaMP6s infection of iSPNs in the VLS of A2A-cre mice around the implanted optic fiber, as well as VLS iSPN axons located under the fiber in the ventral part of the GPe (Fig. 1B), the region most densely innervated by the VLS^13,14,15^. We recorded calcium activity as mice behaved in environments geared to encourage the expression of orofacial actions, including food/object licking, biting, and body-licking during self-grooming (see methods). We then isolated all the peaks from these photometry recordings to look at coincident activity between the striatum and axonal regions. Should photometry peaks in the striatum report neuronal firing, we would expect to observe coincident activity in both the striatal and axonal measurements^16,17^.

**Figure 1:**
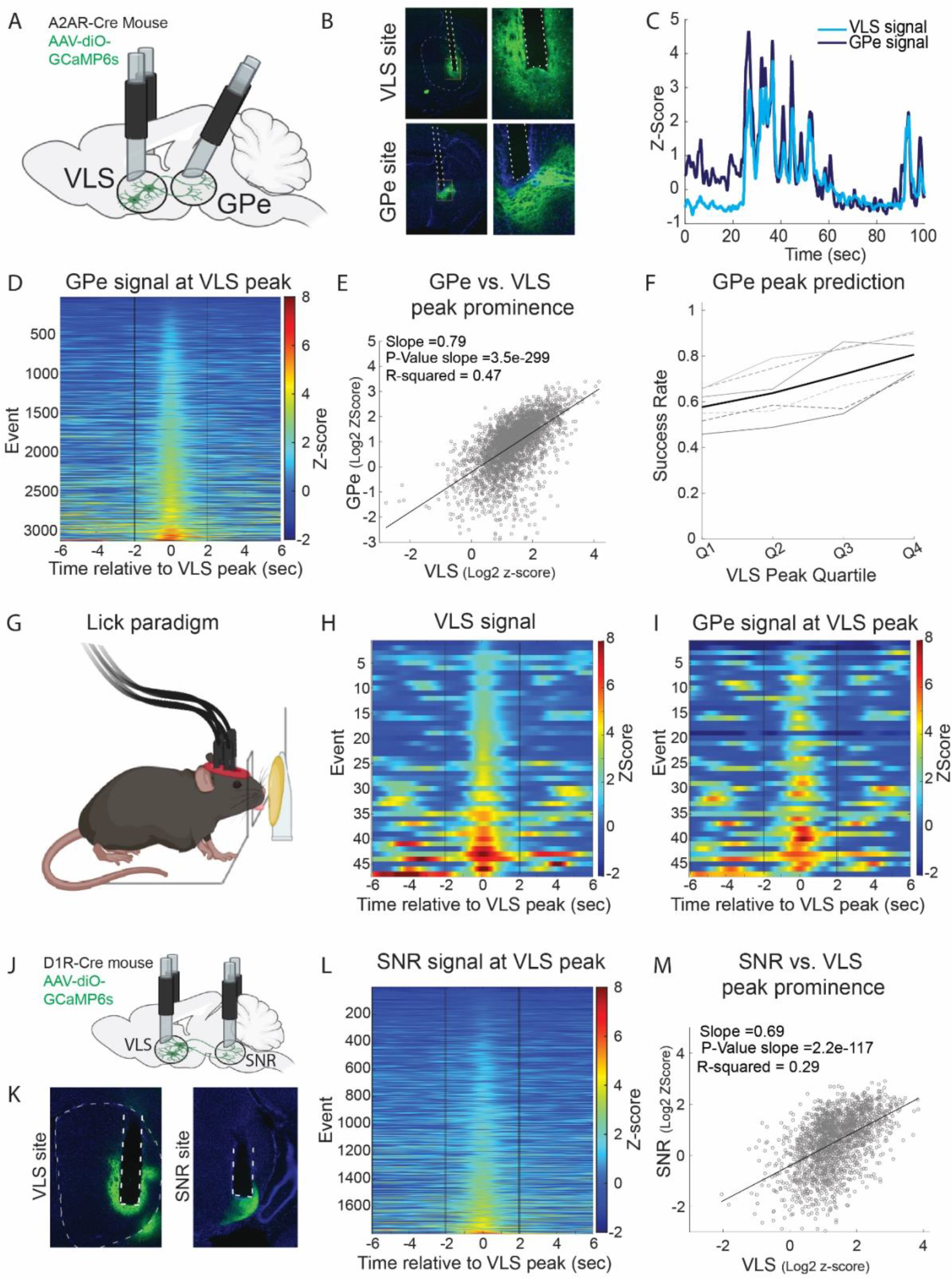
Striatal calcium transients detected by fiber photometry propagate to axons. **A**) Experimental setup for simultaneous recording of fiber photometry signals from soma in the VLS and axons in the GPe of A2AR-Cre mice injected with diO-GCaMP6s in VLS. **B**) GCaMP6s infection and fiber optic cannula placement in VLS vs GPe of representative mouse. White dotted lines in both panels indicate cannula implantation site. In striatal section, striatal boundary is indicated by thinner dotted lines. **C**) Example 100-sec time epoch depicting simultaneous recording of VLS and ipsilateral GPe photometry signals. **D**) Heatmap depicting the Z-score traces of GPe signal recorded during events detected in the VLS signal (z-score> 2; sorted by maximum VLS z-score). Global correlation across all recordings R=0.78, while around peak R=0.86 (-2 to +2 sec relative to peak). N=3123 events from 6 hemispheres from 3 mice. **E**) Plot of log2 maximum z-score of VLS signal plotted against log2 maximum z-score of simultaneous GPe signal for each VLS peak event (local prominence > 2). 3123 events from 6 hemispheres from 3 mice. Slope, p-value, and r-squared value of corresponding fit are defined in the panel. **F**) Success rate in predicting a GPe peak (z-score>2) coincident with a VLS peak, quartiled by VLS peak-prominence (min. z-score prominence >= 2.5, n = 2241 peaks). **G**) Schematic of self-initiated butter-licking paradigm. Licks were scored by the crossing of the threshold by the mouse’s tongue (see methods). **H** and **I**) Side-by-side representation of z-score traces in VLS (**H**) and GPe (**I**) across VLS-peaks coincident with scored licks (from 6 hemispheres across 3 mice). Correlation between signals around peak R=0.89 (-2 to +2 sec relative to peak). **J**) Schematic illustrating dSPN soma and axon recording setup in Drd1-Cre mice. **K**) GCaMP6 infection and fiber optic cannula placement in VLS and SNR of representative mouse. White dotted lines in both panels indicate cannula implantation site. In striatal section, striatal boundary is indicated by thinner dotted lines. **L**) SNr z-score traces recorded simultaneous to VLS-peaks (z-score > 2) as in F, sorted by maximum VLS z-score. Right hemispheres from 3 mice. Global correlation across all recordings = 0.53, while correlations at peaks R=0.68 (-2 to +2 sec relative to peak). **M**) Plot of log2 maximum z-score of VLS signal plotted against log2 maximum z-score of simultaneous SNr signal for each VLS peak event across n = 3 mice, as in F. 1800 events from 3 mice. Slope, p-value, and r-squared value of corresponding fit are defined in the panel.

Visualizing a representative trace of Ca^2+^ activity in both VLS iSPNs and in their axons in the GPe, we observed coincidence at VLS peaks (Fig. 1C). We next isolated peaks of VLS signal from all our recorded trials (z-score>2; n=6 hemispheres from 3 mice) and visualized the coincident axonal signal in the ipsilateral GPe. As can be seen in the corresponding heatmap (Fig. 1D), the vast majority of VLS events were associated with a coincident peak in the GPe signal. There were very few events in which no coincident increase in GPe activity was observed (Fig. 1D). Comparing the prominence, or magnitude, of the peaks for each of the individual events we further observed a strong correlation (Fig. 1E). Notably, individual instances in which VLS and GPe event z-scores were uncorrelated do exist, albeit being a minority of events (Fig. 1D, Fig. 1E). These events, in which there was a VLS peak and yet a coincident GPe peak was not observed could represent occasions in which the striatal calcium signal does not propagate to the axons, and may relate to neuropil activity. We next addressed the reliability of signal propagation from the soma/dendrite area to the axons (defining a lower threshold of z-score>2 for a detected event in the axonal signal). As would be anticipated from a correlated signal, we observed that the larger the signal in the VLS, the greater the probability of observing a corresponding significant peak in downstream axons in the GPe (Fig. 1F).

Ultimately, the primary interest of the field lies in addressing the relationship between neural activity and behavior. The VLS has already been shown by several independent approaches to be important for executing goal-directed orofacial behavior^13,18,19^. Therefore, we addressed the relation of VLS signal to axonal activity during self-initiated licking behavior in freely-moving mice. To this end we established a paradigm in which food-restricted mice licked for a butter reward through a slit in the open-field arena (Fig. 1G). VLS iSPN peaks that overlapped with the initiation of licks were isolated, and the corresponding VLS iSPN photometry signal was plotted (Fig. 1H), alongside the coincident signal from the ipsilateral GPe (Fig. 1I). These heatmaps demonstrate a tight coupling of the GPe axonal signal to the VLS iSPN somatic signal in the context of relevant behaviors. These results illustrate that the vast majority of events recorded directly from a population of behaviorally-relevant striatal neurons propagate in a highly reliable fashion to the axons of these neurons.

We wanted to confirm that the degree of correlation between striatal and axonal signals hold true for dSPNs as well. Employing a similar strategy, recording from VLS dSPNs and axons in the SNr in Drd1-CRE mice (Fig. 1J, Fig. 1K), we observed peaks in the signal recorded in the ipsilateral SNr that were aligned to VLS signal peaks (Fig. 1L). We also observed a correlation between the size of the peaks observed in the VLS and those recorded from axons resident in the SNr (Fig. 1M).

Taken together, our findings suggest that while the transformation from activity in the soma region to axonal activity is not perfect, Ca^2+^ signal peaks recorded in the soma region correlate strongly with axonal Ca^2+^ signal peaks. In particular, the largest of the peaks in the striatum predict with very high fidelity that neural activity will be transmitted from the soma along the axon. While our data do not provide any direct, spatially-precise evidence as to the intracellular source of the Ca2+ signal, our observations support the interpretation of calcium transients in the photometry signal as representing neural activity which propagates to axons in the majority of instances.

In this study, we provide a methodology to measure the axonal transmission of neuronal calcium photometry events, through coincident neuronal and axonal measurement. Importantly, the correlation in signal between the neuronal- and axonal-recording fibers we observed is very likely an underestimate of the true correlation between these two subcellular compartments. This is because it is virtually impossible to record selectively from the exact same population of neurons and their axons due to technical limitations. In actuality, the GCaMP6s+ axons recorded with the GPe/SNR fiber may only partially overlap with the axons whose soma-regions were recorded by the VLS fiber, or likewise may include GCaMP^+^ axonal fibers stemming from neurons that were peripheral to the light path of the fiber located in the striatum (Fig. S1). Thus, the true neuronal-axonal signal correlation is likely even higher than that which is reported here. Looking forward, other researchers may find this experimental paradigm useful for validating the propagation of photometry signal in their neural population of interest. Our strategy can be additionally employed for investigating the variability of signal propagation from somato-dendritic regions to axons *in vivo*, which may reflect biologically interesting integration during behavior^8^. In summary, our observations support the interpretation of calcium transients detected with fiber photometry as reliably reporting action potential activity which propagates to axons.

## Methods

Stereotactic injection and cannula implantation: iSPN mice: A2AR-cre mice (∼14 weeks old; n = 3) were bilaterally injected with 75nL AAV9-CAG-diO-GCaMP6s (titer = 3.0 x 10^12 vg/ml) to the VLS (x = +/-2.5, y=+0.4, z = -4.0) at a 16-degree posterior angle. Neurophotometrics cannulas of diameter 400um were then implanted in the VLS (x = +/-2.5, y=+0.4, z = -3.9), and in the GPe (x = +/- 2.12, y=-0.65, z=-4.25). Cannulas were implanted at a 10-degree posterior angle so that tethering of posteriorly-implanted axonal-recording cannulas would not interfere with the tethering of striatally implanted fibers. dSPN mice: n = 3 D1-cre mice (∼14 weeks old) were similarly injected with 75nL AAV9-CAG-diO-GCaMP6s (titer = 3.0 x 10^12 vg/ml) to the VLS (x = +/-2.5, y=+0.4, z = -4.0) at a 16-degree posterior angle and 400um Neurophotometrics cannulas were implanted vertically in both the VLS (x = +/-2.5, y=+0.4, z = -3.9), and in the SNR at (x = +/-1.5, y=-3.52, z = -4.05). All coordinates are relative to bregma.

Fiber photometry: Recordings were made using the Neurophotometrics fiber photometry setup, where blue photometry-signal wavelength light (470 nm) is time-multiplexed with isosbestic-recording wavelength light (415 nm) at a user-defined frame rate of 30 fps to complete one cycle of both wavelengths of light at 15 Hz. Light power at the fiber tip was set between 100-200 uW for all experiments performed.

Behavior: Mice, aged 3-6 months, were subjected to one 40-minute photometry trial per day in an empty arena composed of plexiglass floor and walls. Trials were aimed to encourage orofacial action, and comprised of either: A) Exposure to the arena with no stimulus; B) 10% sucrose splash on the backside of the mouse to elicit body licking; C) 300mg butter spread on the floor, and D) 300mg butter placed just outside a slit opening in the arena wall. The night before butter trials, mice were rationed 1g food per mouse to induce a moderate hunger state the following day. Cameras (Microsoft Lifecam 3000) were positioned outside the arena to capture behavioral data. For butter-slit trials, one camera was positioned orthogonal to the mouse’s head when it poked through the slit, capturing extensions of the nose and tongue through the slit.

Data Analysis: Recordings were analyzed using custom code in Matlab. Both 470nm signal and isosbestic (415nm) signals were individually smoothed using a local linear regression filter (‘lowess’ filter in Matlab) to reduce high-frequency noise, as in^20^. The slope of the isosbestic curve was then calculated using Matlab’s polyfit function with a degree of 1, and the isosbestic signal was curve-fit to the signal channel^20^. The fitted isosbestic recording trace was subtracted from the raw signal trace and deltaF/F was calculated, and then transformed to z-scores across the whole 40-minute recording timeline. Recordings in which no significant signal was detected (i.e. the z-score did not fluctuate above or below 0 over 10 consecutive minutes; 2 out of 25 recording sessions for the iSPN dataset) were removed from analysis. For the dSPN data-set only recordings from the right hemisphere were included, since we experienced technical issues with the implantation of the left cannula. Several initial filtering conditions were examined, such as using a zero-phase moving average-filter (filtfilt in Matlab)^21^. Results were robust across filtering conditions, and correlated with signal changes in the signal (GCaMP) channel, and not the isosbestic channel.

Signal and Peak Analysis: Peaks were isolated using Matlab’s peak-finder function, with each peak defined as having a minimum-peak prominence of at least 2. Peak tables were then created containing the z-scores of 12-second intervals around the target site (VLS-ipsilateral) defined peaks, and signals were analyzed both in the VLS and GPe/SNR sites at these peak-isolated time-points. For panels 1G and 1L, all raw signal traces from axonal regions were sorted according to max GPe/SNR z-score and subsequently plotted. For correlation plots in panels 1F and 1K, all maximum z-score calculations were performed on z-score traces across events after first subtracting a baseline z-score value for the z-score event trace from each site – the baseline was defined as the average z-score of the period -6 to -4 seconds before the VLS peak. In panel 1H, the success rate was calculated as the percentage of VLS z-score peaks in each quartile that overlapped with a GPe peak of z-score 2 or greater. For 1I and 1J, VLS peaks overlapping with behaviorally relevant events were defined in the following manner: First, VLS-peak tables were assembled as described (12-second event traces surrounding each site of VLS peak - prominence > 2). Then, a peak was defined as coinciding with a lick through the butter-slit if it occurred coincidently with a scored lick. A lick was defined as the bottom half of the most anterior part of the mouse (almost always corresponding to the mouse’s tongue) crossing a threshold ROI line drawn at the border of the butter smear, as captured by a side camera positioned orthogonal to slit-opening in the arena sidewall. An example movie snippet showing scored licks across a whole recording session is provided in supplementary files. Then, for each VLS-peak/lick event, a baseline-subtracted z-score trace (baseline = average z-score -6 to – 4 sec relative to peak, as defined previously) was computed for each site. Finally, the resultant simultaneous baseline-subtracted z-score traces were plotted for each site.

## Author Contributions

DML conceived the approach, performed the majority of experiments and analyses and wrote the manuscript. MT contributed to surgeries and data collection. IS, TS & BJG contributed to the conceptual and technical development of the project, including experiments not included in the final version. MG produced AAV viruses. AC obtained funding, supervised the project and wrote the manuscript.

---

**Supplementary Figure 1:**
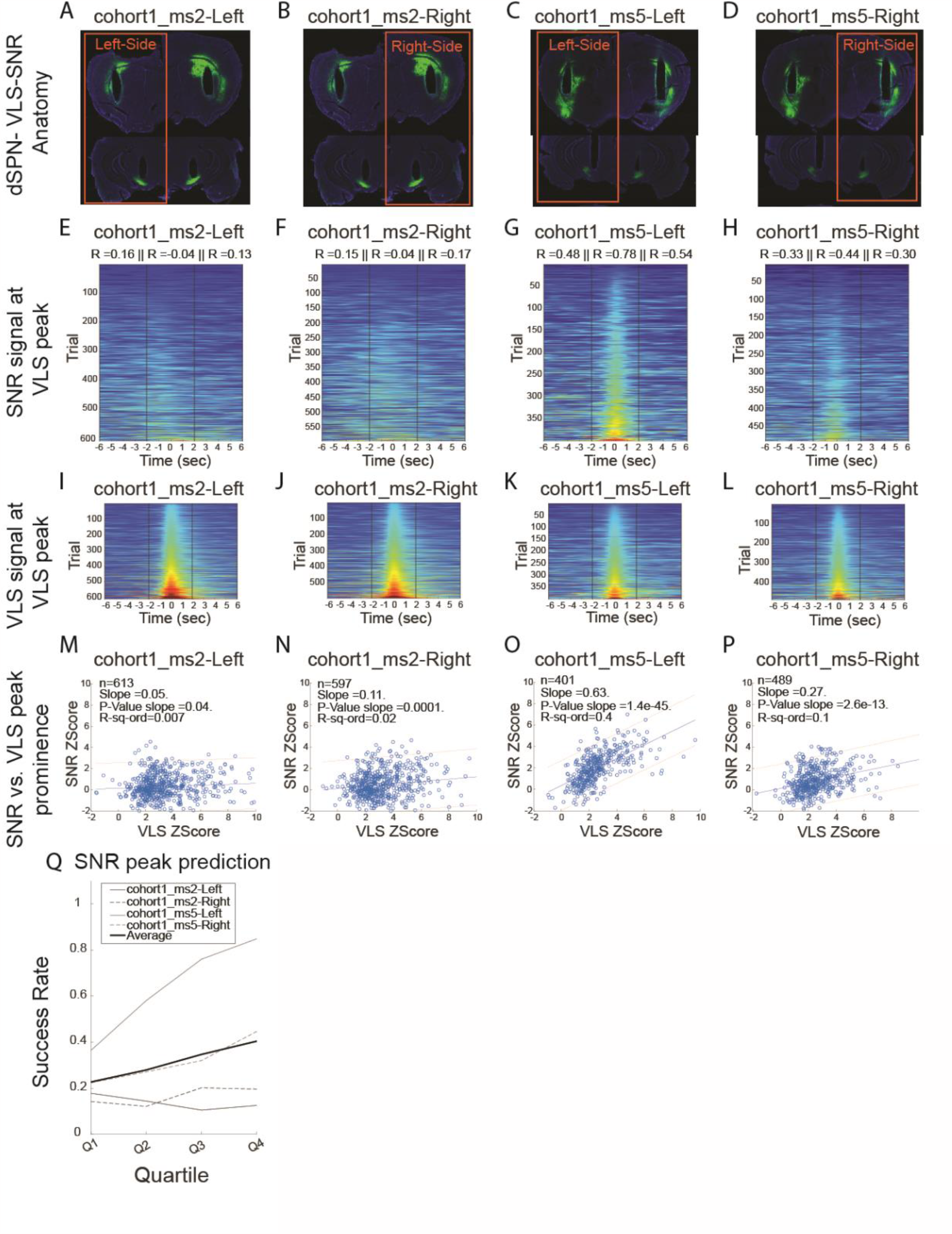
Analysis of Striatal/SNR mismatch vs. on-target injections. A-D) Striatal and SNR sections showing GCaMP6s+ infection of neurons in striatum and GCaMP6s+ axons in SNR for each hemisphere of two mice from dSPN VLS-SNR cohort-1, ms#2 and ms#5. In ail mice, the striatal fiber is placed consistently above GCaMP6s+ neurons in the VLS and measures activity of VLS dSPNs. Mice exhibit variability in the extent of infection of VLS versus DLS/CTX GCaMP6s+ neurons, and therefore varying contribution of DLS/CTX versus VLS SPN axons to the signal from the SNr fiber. E-H) Heatmaps of SNr signal at VLS peak for each mouse hemisphere whose anatomy is shown in A-D, similar to Figure 1L. Peaks sorted by max SNR Z-score. Corrélation values are shown for 4-second intervals around VLS peak: i) -6 to -2 sec prior to peak, ii) -2 to +2 sec around VLS peak. iii) 2 to 6 sec after sig al peak. I-L) Heatmaps of VLS signal at VLS peaks for same data set of peaks as shown in E-H, sorted by VLS max. Z-Score. M-P) Plot of maximum z-score of SNR-ipsilateral signal plotted against maximum z-score of VLS-ipsilateral signal at VLS-ipsilateral peaks, similar to Fig. 1M for each mouse hemisphere as in A-D. Q) Success rate of peak prédiction for cohort1 mice.

